# Identification and quantification of chimeric sequencing reads in a highly multiplexed RAD-seq protocol

**DOI:** 10.1101/2021.09.21.461194

**Authors:** Maria Luisa Martin Cerezo, Rohan Raval, Bernardo de Haro Reyes, Marek Kucka, Frank Yingguang Chan, Jarosław Bryk

**Affiliations:** Department of Biological and Geographical Sciences, School of Applied Sciences, University of Huddersfield, Queensgate, Huddersfield, England, United Kingdom; AVIAN Behavioural Genomics and Physiology, IFM Biology (IFM), Linköping University, Linköping, Sweden; Friedrich Meschier Laboratory of the Max Planck Society, Tübingen, Germany

**Keywords:** Chimeras, RAD-seq, quaddRAD, index hopping, read misassignment, adapters, barcodes

## Abstract

Highly multiplexed approaches have become a common practice in genomic studies. They have improved the cost-effectiveness of genotyping hundreds of individuals by using combinatorially-barcoded adapters. These strategies, however, can potentially misassign reads to incorrect samples. Here we used a modified quaddRAD protocol to analyse the occurrence of index hopping and PCR chimeras in a series of experiments with up to a 100 multiplexed samples per sequencing lane (total n = 639). We created two types of sequencing libraries: four libraries of Type A, where PCR reactions were run on individual samples before multiplexing, and three libraries of Type B, where PCRs were run on pooled samples. We used fixed pairs of inner barcodes to identify chimeric reads. Type B libraries show a higher percentage of misassigned reads (1.15%) compared to Type A libraries (0.65%). We also quantify the commonly undetectable chimeric sequences that occur whenever multiplexed groups of samples with different outer barcodes are sequenced together on a single flow cell. Our results suggest that these types of chimeric sequences represent up to 1.56% and 1.29% of reads in Type A and B libraries, respectively. We review the source of such errors, provide recommendations for developing highly-multiplexed RAD-seq protocols and analysing the resulting data to minimise the generation of chimeric sequences, allow their quantification, and provide finer control over the number of PCR cycles necessary to generate enough input DNA for library preparation.

## 1 Introduction

Development of high-throughput sequencing and reduced representation approaches, such as restriction-site–associated DNA sequencing (RAD-seq), have dramatically reduced the cost of generating vast amounts of sequencing data. RAD-seq (Baird *et al*., 2008) and its many variants, including but not limited to: ddRADseq (Peterson *et al*., 2012), quaddRAD-seq (Franchini *et al*., 2017) and adapterama (Glenn *et al*., 2019; Bayona-Vásquez *et al*., 2019) have been used in studies of phylogenetics (Massatti *et al*., 2016; Lecaudey *et al*., 2018; Near *et al*., 2018), phylogeography (Jeffries *et al*., 2016), association mapping (Nadeau *et al*., 2014), introgression (Hohenlohe *et al*., 2013), population structure and genetic diversity (Rodríguez-Ezpeleta *et al*., 2016; Leone *et al*., 2019; Gao *et al*., 2017; Martin Cerezo *et al*., 2020).

Many high-throughput methods rely on multiplexing, inclusion of unique identifying sequences in the adapters of each sample (barcodes or indices), pooling of samples, and subsequently sequencing pools on a single sequencing lane. This approach has now become common practice, and single (Poland and Rife, 2012), dual (Glenn *et al*., 2019; Peterson *et al*., 2012) and quadruple barcodes (Franchini *et al*., 2017; Bayona-Vásquez *et al*., 2019) have been developed. While these approaches greatly reduce the costs of sequencing, they can also increase the number of misassigned reads (MacConaill *et al*., 2018), where sequences from one sample are incorrectly assigned to another due to sequencing errors, nucleotide misincorporations and contamination of adapters during synthesis or library preparation (Van Der Valk *et al*., 2019), amongst others.

PCR chimeras are reads composed of distinct parental sequences, and are one source of read misidentification (Fonseca *et al*., 2012). Spontaneous dissociation of polymerases from the template molecules during amplification can occur due to low processivity, secondary structures of DNA, or incomplete extension of DNA during the PCR (Smyth *et al*., 2010), and can lead to the formation of incomplete sequences. These fragments can act as primers during subsequent amplification cycles, and produce artificial PCR products containing fragments of sequences containing barcodes from two different samples. Similarly, index hopping, caused by free-floating primers resulting from insufficient DNA purification, erroneous size selection of the library, or due to fragmentation during improper storage, can prime template DNA molecules on a sequencing flow cell prior to exclusion amplification (Van Der Valk *et al*., 2019). The presence of both chimeric and index-hopped sequences can lead to inflated measures of diversity, and bias population genetics parameters in downstream analyses (Smyth *et al*., 2010; Van Der Valk *et al*., 2019).

Additionally, chimeras can occur whenever a single flow cell is filled with groups of samples that were processed independently but which share inner barcodes, such as in quaddRAD (Franchini *et al*., 2017). In this case, the chimeric sequences are impossible to differentiate from genuine samples during downstream analysis. Only when unique combinations of both inner and outer barcoded adapters are used, can such chimeric reads be identified, quantified and eliminated during analysis. While PCR-free protocols for library preparation can alleviate the problem of read misassignment, studies have found considerable read misassignment in these libraries (Costello *et al*., 2018). Furthermore, they require a large amount of high-quality DNA, which is often not available.

The incidence of index-hopping during cluster generation of the Illumina HiSeq-X and NovaSeq platforms has been reported as less than 1% (Van Der Valk *et al*., 2019). However, the still widely used platforms utilising exclusion amplification (ExAmp) cluster generation such as the HiSeq 3000/4000 have reported misassignment rates up to 10% (Sinha *et al*., 2017), and up to 30% in PCR reactions (Wang and Wang, 1996). We note, however, that published analyses of chimeric sequences were obtained on relatively few samples, and considered formation of chimeric sequences only during cluster generation or only during library preparation.

Here, we quantify the prevalence of chimeric sequences in two large-scale, highly multiplexed experiments (86 to 100 samples multiplexed per lane of sequencing, total number of samples = 639). We assessed the contribution of both PCR amplification and sequencing to the generation of chimeric sequences, by preparing two types of libraries: Type A, where adapter ligation and PCR amplification was carried out on each sample individually, and Type B, where samples were pooled before amplification and addition of outer adapters, respectively. Our design also allows for identification of chimeric sequences that are otherwise undetectable: sequences formed between groups of samples that share some combinations of inner adapters, but that were processed in different multiplexed groups (Figure 2, Step 2). Overall, we identify and quantify four types of chimeric sequences (Type I - IV, Figure 2, Step 3 and Methods 2.4). The details of the experimental design and illustration of the different types of chimeras are shown in Figure 2. Based on our findings, we provide recommendations for adapter design and data analysis to minimise the number of misassigned reads.

## 2 Material and Methods

### 2.1 Adapter design

Adapters were designed following the quaddRAD protocol (Franchini *et al*., 2017). Restriction enzyme overhangs were modified for SbfI and MseI. 8bp-long barcodes were designed using EDITTAG (Faircloth and Glenn, 2012), with a minimum Levenshtein distance of 4 nucleotides, GC content of 40-60% and avoiding sequences that were self-complementary and containing more than two adjacent, identical bases. From 102 tags suggested by EDITTAG, sequences that reconstructed SbfI and MseI restriction sites were removed manually. 18 tags were selected for the inner adapters and 8 for the outer adapters, giving a total of 144 possible combinations.

Four random nucleotides (5’-VBBN-3’) were also incorporated into the inner adapters to allow *in silico* identification of PCR duplicates (Figure 1). Inner adapters were used in fixed pairs while outer adapters have been used combinatorially. A complete list of adapter sequences can be found in Supplementary Materials Tables 1-3.

**Figure 1:**
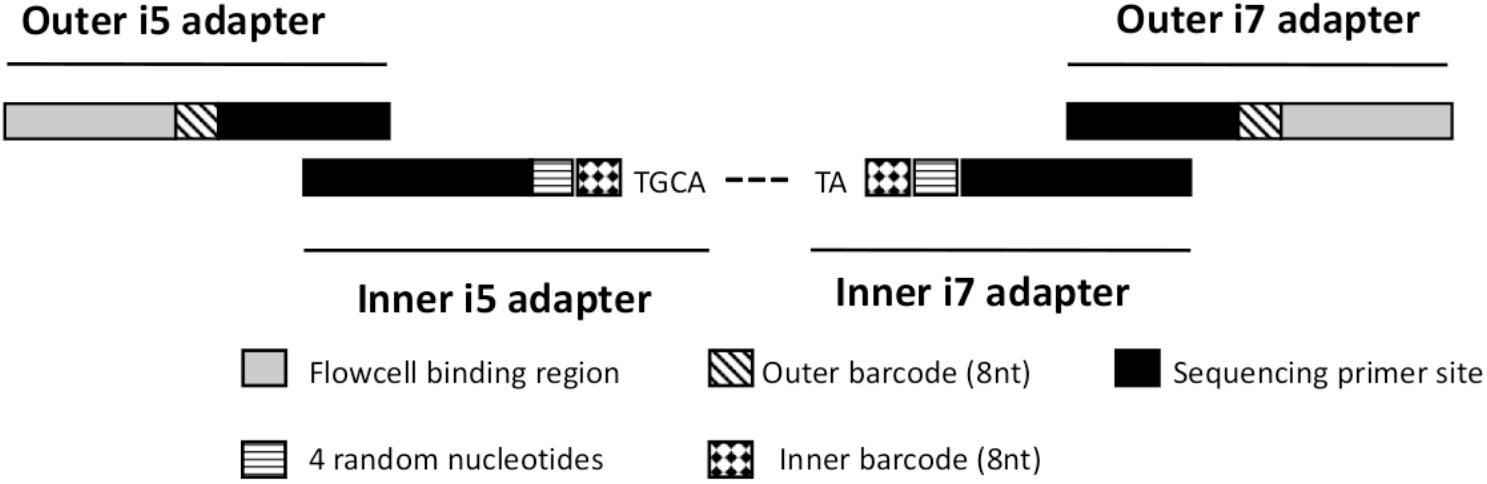
Elements of adapter sequences. Modified from Franchini *et al*. (2017)

### 2.2 Library preparation

DNA from 459 *Apodemus flavicollis* and 180 *Apodemus sylvaticus* tissue samples were extracted following Martin Cerezo *et al*. (2020). Seven libraries were prepared following a modified version of Franchini *et al*. (2017) protocol.

Four libraries (n = 164 *A. flavicollis* and 180 *A*.*sylvaticus*, 86 samples per library), henceforth called Type A, were prepared with each sample individually amplified to allow for the quantification of sequencing chimeras or index-hopped reads only. The three remaining libraries (n = 295 *A. flavicollis* samples including 96, 99 and 100 samples per library), henceforth called Type B, were multiplexed following restriction digestion and ligation of barcoded inner adapters before they were amplified as a pool. This allowed for the quantification of the total number of chimeric sequences, which originated both during sequencing and during PCR amplification.

#### 2.2.1 Type A libraries: individual PCR reactions

Inner adapters were prepared by annealing each single-stranded oligonucleotide with its complementary strand. 5 µl of each bottom and top strands at 100µM were mixed with 40 µl of annealing buffer (50 mM NaCl, 10 mM Tris-Cl, pH 8.0), heated to 98°C for 2.5 min and cooled at a rate of 1°C per minute down to 15°C. Once prepared, the adapters were kept at -20°C and used within 2 weeks.

60 ng of genomic DNA was then digested and ligated to the inner adapters in a single-step 40 µl reaction containing 4 µL 10x CutSmart buffer, 1.5 µL *Mse1* (10 U/µL), 0.75 µL *Sbf1* (20 U/µL), 4 µL ATP (10mM), 1 µL T4 DNA ligase (400 U/µL), 0.75 µL of each quaddRAD_i5n and quaddRAD_i7n inner adapters (10 µM), ddH_2_O to 40 µL and incubated for three hours at 30 °C in a thermocycler. The reaction was stopped with 10 µl of 50 mM EDTA. Samples were purified and double size selected using 0.4x and 0.8x Sera-Mag SpeedBeads solution (GElifesciences, Marlborough, MA, USA) containing 10 mM Tris base, 1 mM EDTA, 2.5 M NaCl, 20% PEG 8000 and 0.05% Tween 20 (pH 8.0), and eluted in 30 µL 10mM Tris-HCl.

To introduce the outer barcoded adapters, an indexing PCR was carried out in a 50 µl reaction containing 4 µl of each i5 and i7 primers (5mM), 1 µl of dNTPs (10 mM), 10.5 µl of purified water, 10 µl of 5x Q5-HF Buffer, 0.5 µl of Q5-HF DNA Polymerase (New England Biolabs, Frankfurt am Main, Germany) and 20 µl of template DNA. After an initial denaturation step of 30 seconds at 98°C, the PCR reaction was carried out in 14 cycles (15 seconds at 98°C, 30 seconds at 67°C and 60 seconds at 72°C) and a final elongation at 72°C for 2 minutes. Purification was performed using 0.8x Sera-Mag SpeedBeads solution (GElife-sciences, Marlborough, MA, USA) and DNA was eluted in 22 µl Tris-HCl (10 mM).

Samples were multiplexed by combining 10 ng of each sample in Plates 1 and 4 and 20 ng of each samples in Plates 2 and 3. Libraries were then size-selected to 300-600 bp using BluePippin (Sage Science, Beverley, MA, USA) and sequenced on Illumina HiSeq 3000 (Illumina Inc., San Diego, CA, USA).

#### 2.2.2 Type B libraries: multiplexed PCR reaction

Digestion and ligation reactions were performed as described for Type A libraries, except with an initial input of 100 ng of genomic DNA. Samples were then purified using 0.8x SPRI Sera-Mag SpeedBeads solution, eluted in 30 µL of Tris-HCl (10 mM) and subsequently equimolarly pooled according to inner barcode combinations prior to PCR amplification.

An indexing PCR was carried out to introduce the outer barcoded adapters to each pool of digested DNA and enrich the libraries in a 100 µL reaction containing 8 µL dNTP mix (2.5 mM), 20 µL 5x Q5-HF buffer, 4 µL quaddRAD-i5nn primer (10 µM), 4 µL quaddRAD-i7nn primer (10 µM), 1 µL Q5 high-fidelity DNA polymerase (2 U/µL), 50 ng of DNA (restricted, ligated and pooled) and ddH_2_O to 100 µL. Reaction conditions were as described for Type A libraries. Each PCR reaction was again purified using 0.8x SPRI Sera-Mag SpeedBeads solution and eluted in 50 µL of tris-HCl (10 mM). 100 ng of each enriched library were then pooled again and size selected to 300-600 bp using Blue Pippin (Sage Science, Beverly, MA, USA) and sequenced on Illumina HiSeq 3000 (Illumina Inc., San Diego, CA, USA).

### 2.3 Clone removal

Sequences were demultiplexed based on the outer barcodes by the sequencing center (Genome Centre at the Max Planck Institute for Developmental Biology, Tübingen, Germany). PCR duplicates were identified and removed from each library using the clone_filter programme from Stacks 2.41 (Catchen *et al*., 2011) and the random nucleotide tags in the inner adapters. Sequences were then demultiplexed based on the inner barcodes, quality-filtered and truncated to 136bp with process_radtags, also from Stacks, removing reads with uncalled bases, low quality scores, reads that were marked by Illumina’s chastity/purity filter as failing and allowing for barcode and RAD-tags rescue. process_radtags was run 5 times, changing the number of mismatches allowed for barcode rescue from 0 to 4 at each iteration. Samples were demultiplexed using not only combinations of barcodes used for library preparation, but all the possible combinations of inner barcodes, allowing for the quantification of chimeric sequences. The number of retained reads for each barcode combination and for each one of the process_radtags runs were recovered from the log files generated by process_radtags.

### 2.4 Multiplexed groups

multiplexed groups are defined as a set of samples that share outer adapters. Typically, several of such groups are prepared in parallel, pooled, and then sequenced on a single lane of a sequencing machine. Each one of the four Type A libraries included 8 multiplexed groups of 9 samples each, 1 multiplexed group of 8 samples and 1 multiplexed group with 6 samples. Type B libraries had different multiplexed schemes per library. The libraries contained 8 multiplexed groups of 9 samples and 3 multiplexed groups of 8 samples. Library Type B-1 also contained an additional multiplexed group with 4 samples while library Type B-2 contained a multiplexed group with 3 samples.

### 2.5 Identification of chimeric sequences

Based on the combination of inner and outer barcodes used, we divided the identified chimeric sequences into four types. Summary of the experimental design and the types of chimeric reads are shown on Figure 2.

**Figure 2:**
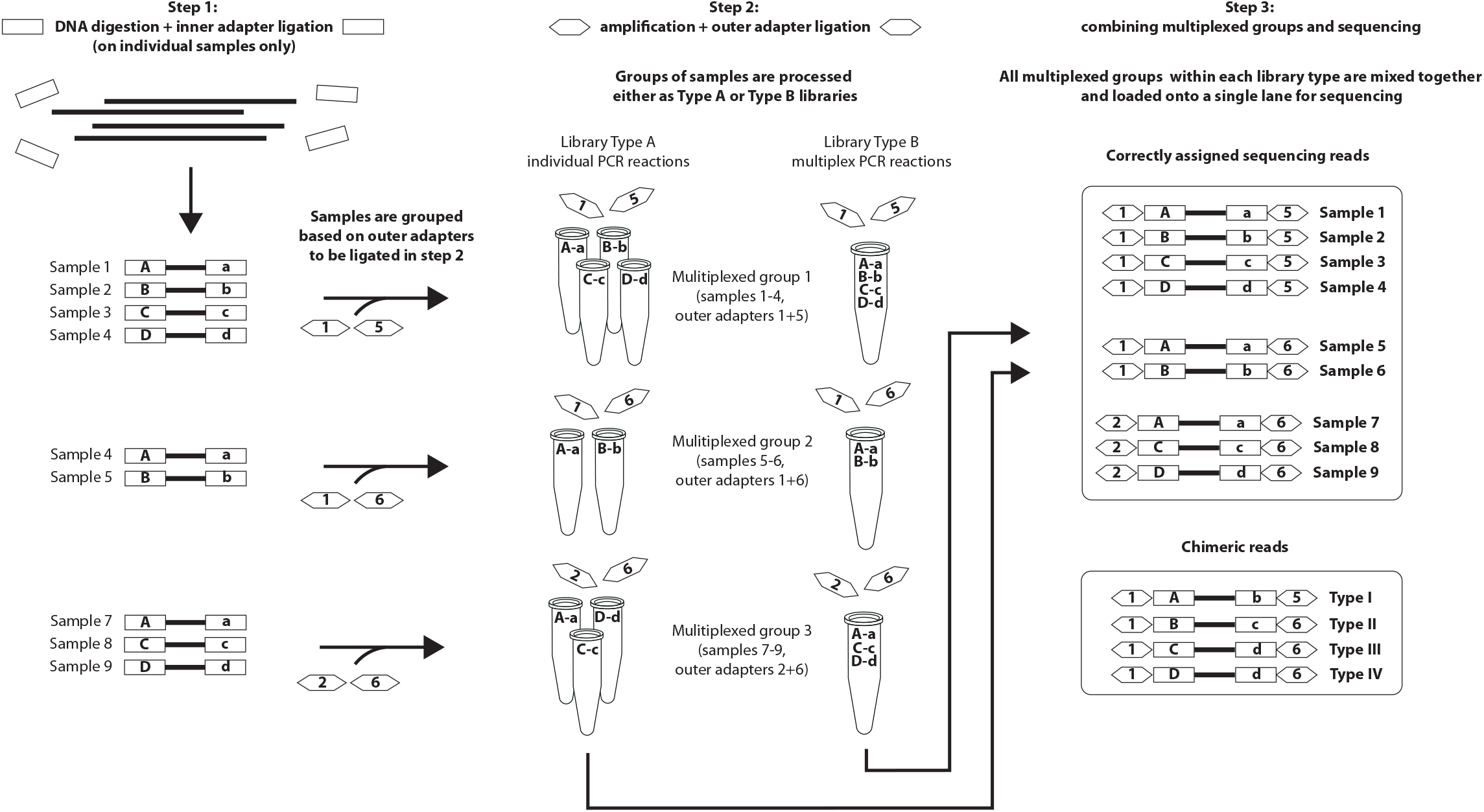
Summary of the experimental design and the types of chimeric reads. Inner adapters are represented by letters and rectangles while outer adapters are represented by numbers and hexagons. Combinations of the same uppercase and lowercase letters represent the unique combinations of inner barcoded adapters used in the protocol. Any other combination of letters represent chimeric combinations of adapters. Multiplexed groups are groups of samples processed together in the protocol.

Type I chimeras are molecules that contain unused combinations of inner barcodes, both of which were used in the same multiplexed group. For example, inner barcodes “A”, “a”, “B” and “b” were present in multiplexed group 1, but not in a combination “Ab”.

Chimeras of Types II-IV form between sequences from different multiplexed groups and are only detectable if multiplexed groups contain different number of combinations of inner barcodes. For example, multiplexed group 1 contains 4 combinations of inner barcodes (Aa, Bb, Cc, Dd) but multiplexed group 2 contains only 2 (Aa, Bb). These chimeras are detectable when samples are processed in several unequally-sized groups but the same set of inner barcodes is used across all groups.

Type II chimeras are reads containing one inner barcode that was used in a different multiplexed group. For example, the combination of inner barcodes “Bc” in multiplexed group 2 is a chimera type II since this multiplexed group contains barcode “B” but not barcode “c”.

Type III chimeras are reads containing a *chimeric* combination of inner barcodes, neither of which was used in their multiplexed group. An example of these type of chimeras is the combination of inner barcodes “Cd” in multiplexed group 2, since this multiplexed group does not contain neither barcodes “C” nor “d”. Barcode “C” was used in multiplexed group 1 and barcode “d” in multiplexed group 3.

Finally, Type IV chimeras are reads containing a correct combination of inner barcodes that were used in other multiplexed groups but not in the group where they were detected. The combination “Dd” in multiplexed group 2 is one of these chimeras since multiplexed group 2 does not contain barcodes “D” nor “d” but this combination of barcodes was used in multiplexed groups 1 and 3. In our protocol, it was only possible to detect Type IV chimeras in 19 out of 75 multiplexed groups as all other groups were equally-sized.

### 2.6 Counts of sequences with chimeric adapters

For each multiplexed group, unused combinations of barcodes were used to detect chimeric sequences and were quantified as a percentage of total sequences within the multiplexed group. Each of the multiplexed groups included between 2 and 9 samples. We calculated percentage of chimeric sequences in relation to mismatches allowed for barcode rescue and the library type. This allowed comparison of the proportion of chimeras when different number of mismatches were allowed, as well as comparison of the results within and between each library type.

To compare the relative abundance of the different types of chimeras, the percentage of chimeric sequences of each type was calculated relative to the number of reads per plate sequenced. Similarly, the percentage of chimeric sequences was also calculated individually for each possible chimeric combination of barcodes.

Type IV chimeras can only be identified in very specific multiplexing schemes and barcodes combinations, but it is worth emphasising that they are being generated in all multiplexed groups, whether or not the experimental protocol enables their detection. We can directly quantify a fraction of Type IV chimeras in 19 of our 75 multiplexed groups, but we expect that in other multiplexed groups or combinations of barcodes Type IV chimeras represent a similar percentage of missassigned reads. We estimated the number of Type IV chimeras, including their generation between all possible combinations of adapters that could produce them, with the following formula:

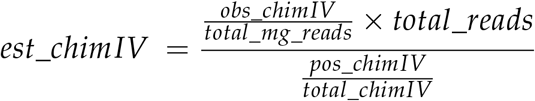

Where:

obs_chimIV: Number of observed type IV chimeras

total_mg_reads: Number of reads in multiplexed groups with observed type IV chimeras

pos_chimIV: Total number of cases that could produce chimeras type IV

total_chimIV: Number of cases where chimeras type IV were identified

total_reads: Total number of reads

Calculations were performed only considering 0 mismatches for barcode rescue.

## 3 Results

### 3.1 Multiplex PCR increases the proportion of sequences with chimeric adapters

Type A libraries, where indexing PCRs were conducted on each sample independently, consistently produced fewer chimeric sequences, as a percentage of total sequences, than Type B libraries, at the same number of mismatches during barcode rescue (Figure 3). Overall, demultiplexing with perfect barcodes showed a median of 0.59% (max = 1.20%, min = 0.33%, mean= 0.65%) and 1.09% (max = 2.33%, min = 0.31%, mean = 1.15%) chimeric sequences for Type A and Type B libraries, respectively.

**Figure 3:**
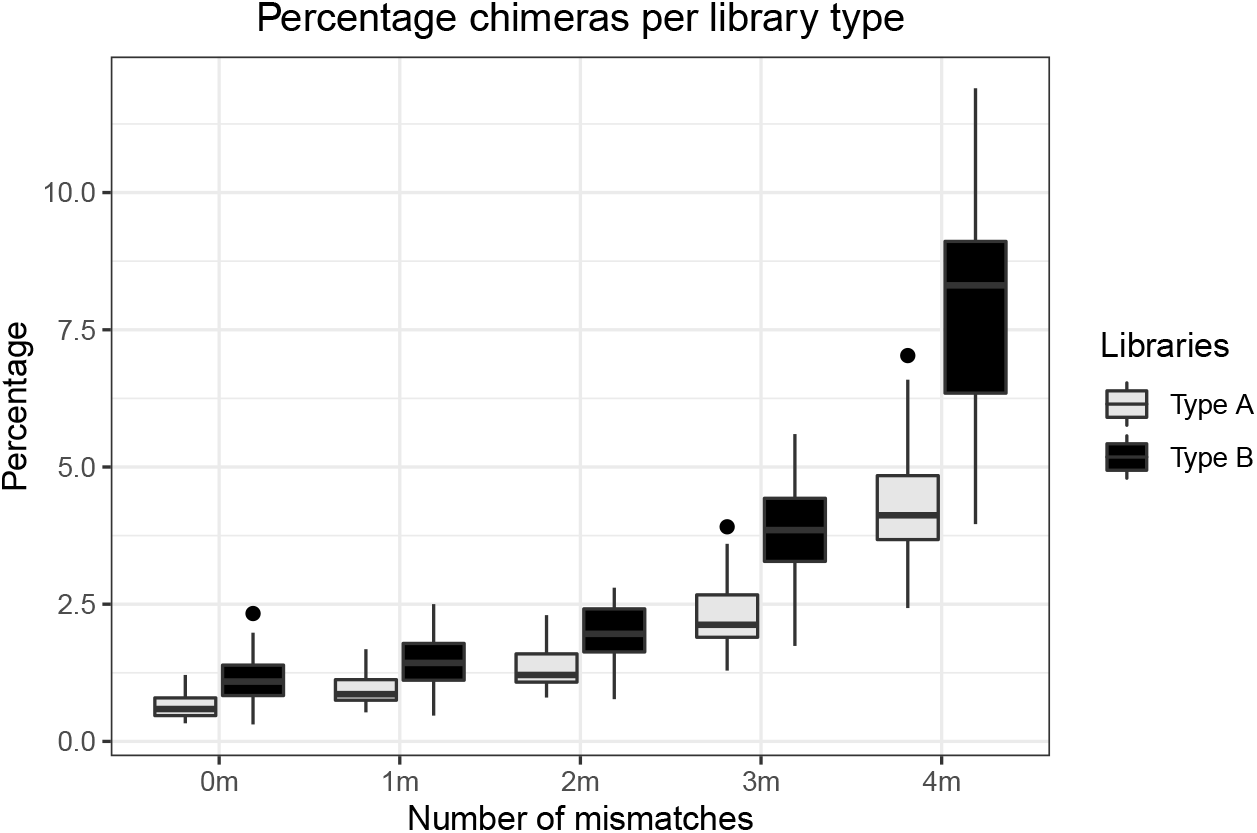
Percentage of chimeric sequences in Type A (grey; PCR on individual samples: includes only sequencing chimeras) and Type B libraries (black, PCR on multiplexed groups of samples, includes PCR and sequencing chimeras) for each level of barcode rescue.

Increasing the number of mismatches for barcode rescue from zero to four also increased the percentage of chimeric sequences detected in both library types to a median of 4.12% (max=7.03, min=2.43, mean = 4.37%) for Type A and 8.31% (max=11.90, min=3.96, mean = 7.85%) for Type B, as a greater number of reads was retained by process_radtags. In all cases, differences between Type A and Type B libraries were significant (Figure 3, Mann Whitney U test, p<0.001).

### 3.2 Differences in percentage of chimeric sequences within Type A or Type B libraries are smaller than between the two libraries

Percentage of chimeric sequences within independently prepared libraries of the same type (four libraries of Type A and three of Type B) were more similar to one another then between libraries of different type (Figure 4). The only differences between libraries of the same type were between Type A-4 vs Type A-1 and Type A-4 vs Type A-3 (*H*_3_ = 13.3, *p* = 0.004, post-hoc Dunn test). No other differences were detected between any other of combination libraries.

**Figure 4:**
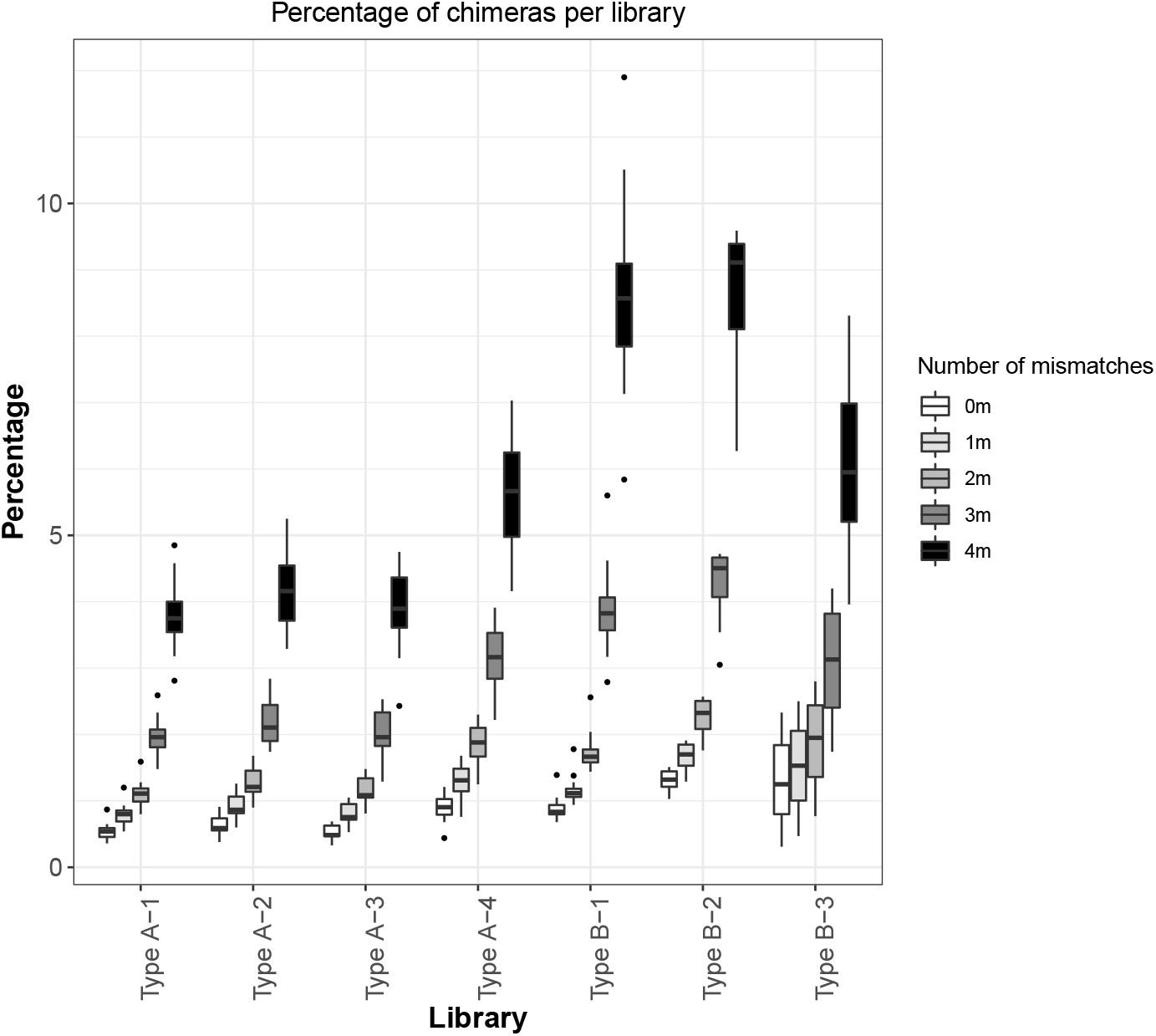
Percentage of chimeric sequences in independently prepared libraries of Type A and Type B for each level of barcode rescue.

### 3.3 The proportion of chimeric sequences increases with the number of mismatches allowed in barcode rescue

Allowing mismatches for barcode rescue enables recovery of sequences with uncalled or erroneous base calls in the barcode sequence. Therefore, any increment in the number of mismatches will increase the number of reads retained after demultiplexing with process_radtags. It will also increase the proportion of chimeric sequences.

The overall proportion of such sequences increased significantly with each increment in the number of mismatches allowed for barcode rescue, from 0 to 4 (Kruskall-Wallis test: *H*_4_ = 287, *p* < 0.001). The closer the number of mismatches is to the distance between barcodes (in our case, 4 nucleotides), the larger the increase in the proportion of chimeric sequences (Figure 5).

**Figure 5:**
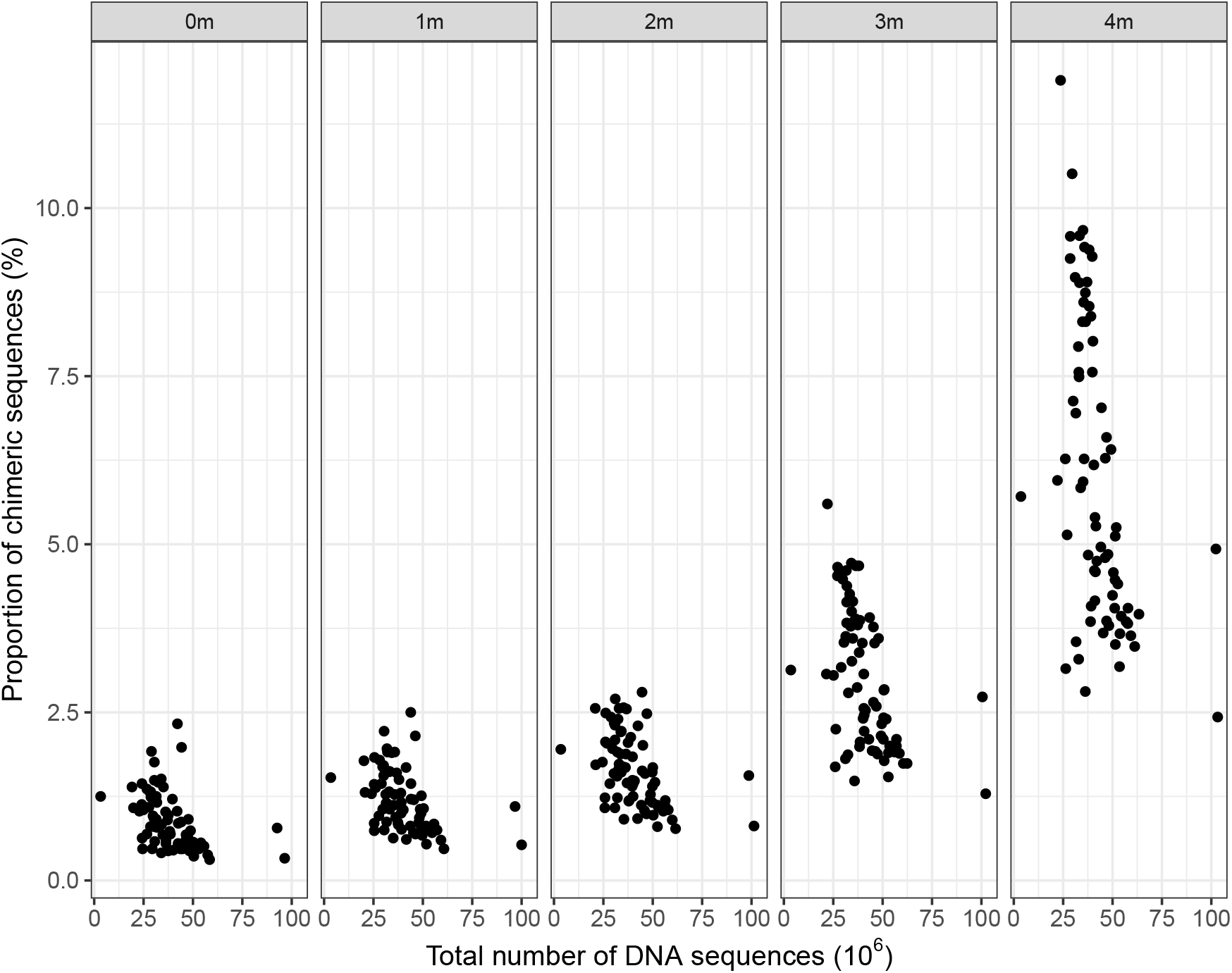
Percentage of chimeric sequences in total number of DNA sequences recovered from both library types for each level of barcode rescue.

Importantly, the number of new reads retained during demultiplexing remain higher than the number of new chimeric sequences detected by process_radtags when we increase the number of mismatches for barcode rescue up to three (Figure 6). Past this point, when the number of mismatches equals the distance between barcodes, the number of new chimeric sequences detected overtakes the number of new retained reads, indicating that increasing the number of mismatches past three has no additional benefit.

**Figure 6:**
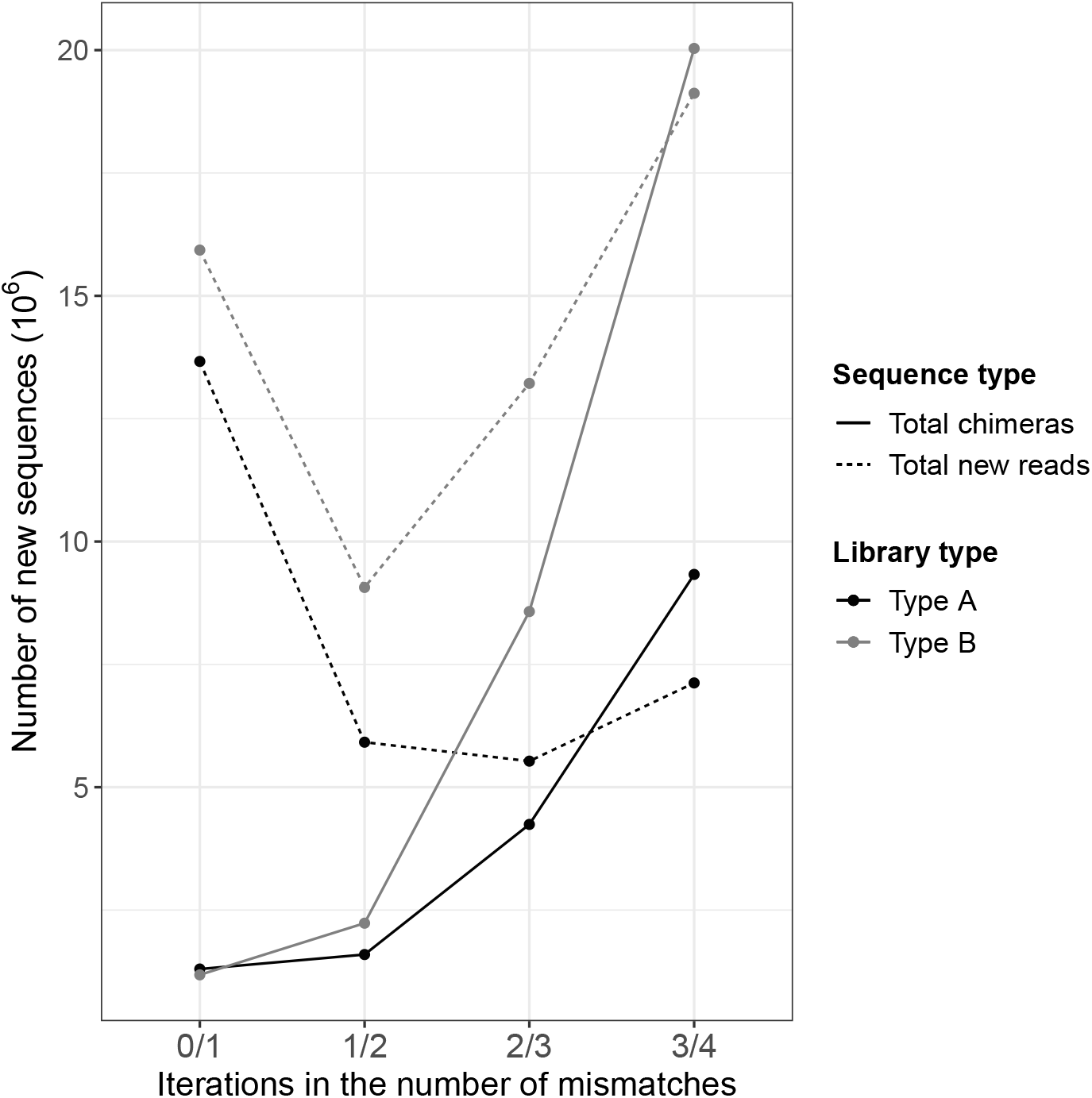
Number of new sequences obtained for each iteration on the number of mismatches allowed for barcode rescue. Solid lines represent the new chimeric sequences while dashed lines represent the total number of new reads. Colour indicates library type (Black: Type A, Grey: Type B)

### 3.4 Quantification of different types of chimeric sequences

When the multiplexed groups are equally-sized, containing 9 samples, among all possible combinations of inner barcodes (n = 81), 11,1% identify genuine reads while the remaining 88,9% identify chimeric sequences. In these cases, 56 out 75 multiplexed groups, only chimeras Type I are detectable. Since chimeras type II-IV are only detectable in a much smaller fraction of multiplexed groups, chimeras type I seem to be the predominant fraction of chimeras per library type (Figure 7). When only chimeras type I are detectable, chimeras type II and III will be identified as chimeras type I while chimeras type IV will be misidentified as genuine samples.

**Figure 7:**
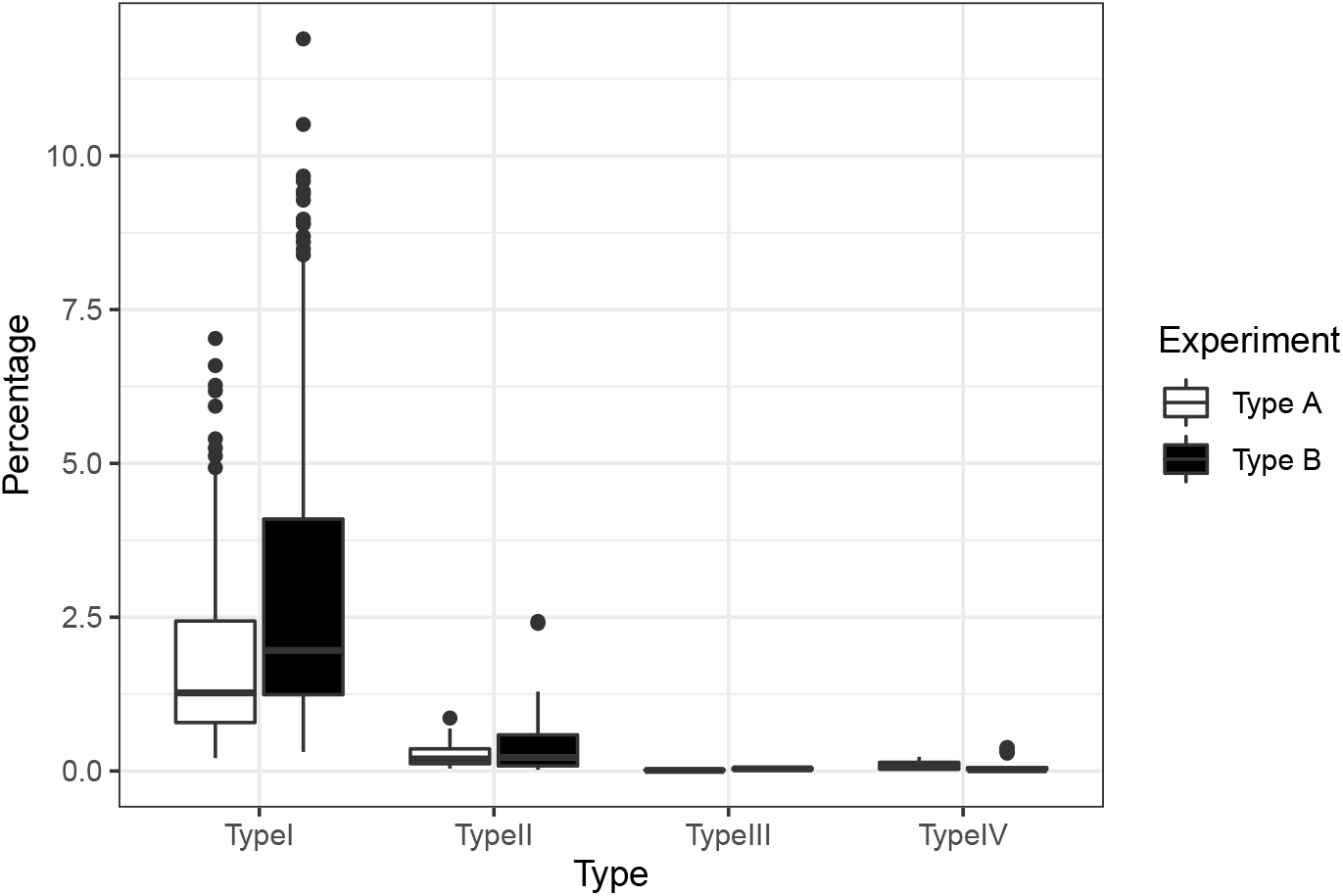
Percentage of chimeric sequences in each library type, relative to the total number of reads per plate sequenced.

In multiplexed groups where the number of multiplexed samples was lower than 9, it is possible to identify chimeras Type II, III and IV. In these cases, 19 out of the 75 multiplexed groups in our protocol, chimeras type IV are the most abundant chimeras when less than 3 mismatches were allowed for barcode rescue (Figure 8).

**Figure 8:**
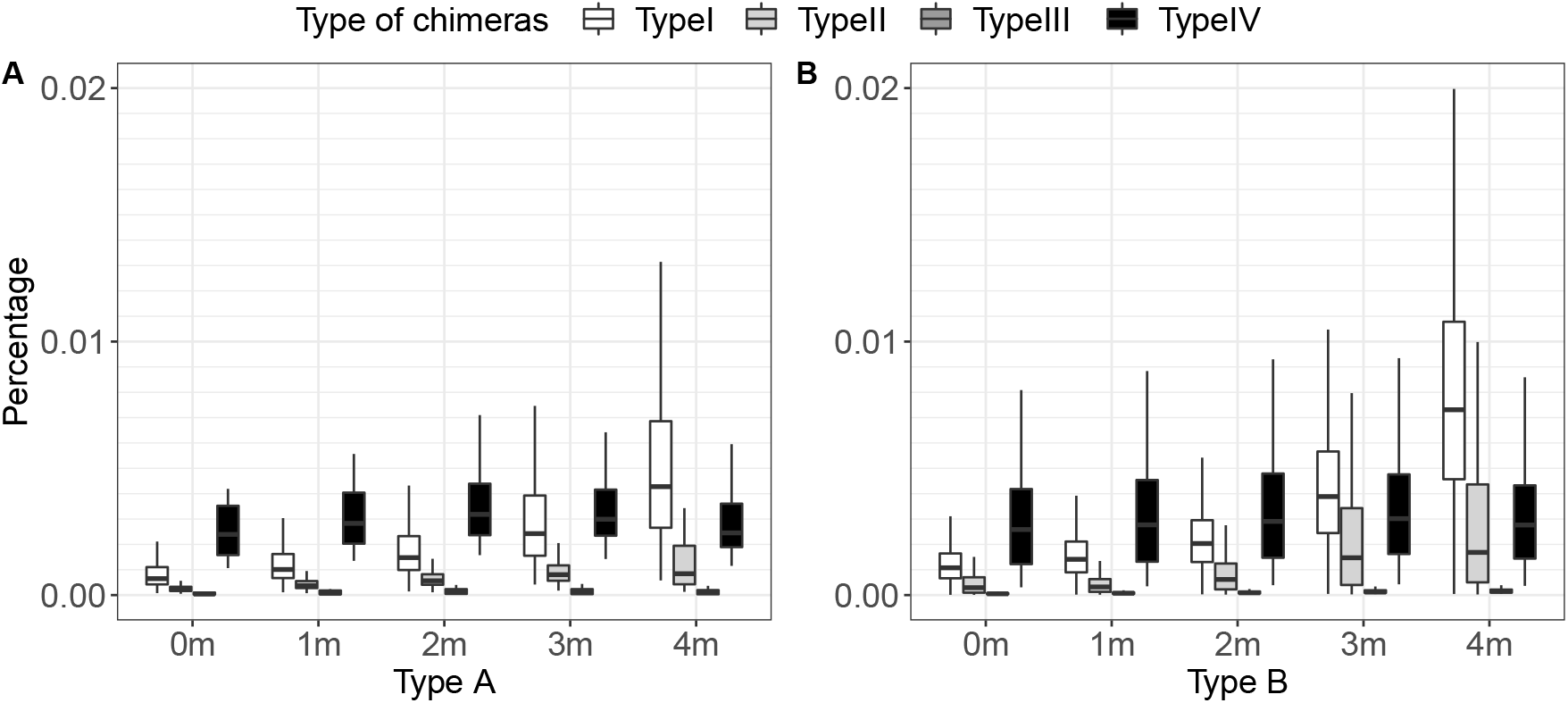
Percentages of four different types of chimeras in each library type. Percentages have been calculated individually for each combination of barcodes.

As it was only possible to detect Type IV chimeras in 19 out of the 75 multiplexed groups, we estimated the expected total number of Type IV chimeras should the protocol had been constructed to detect all the Type IV chimeras. In Type A libraries, 182768 type IV chimeras (0.01% of the total reads from the library) were observed, while the estimated number of type IV chimeras is 28280095 reads (1.56% of the total reads in the library). In Type B libraries, we observed 205518 type IV chimeras (0.019% of the total number of reads in the library) of the estimated 13713202 reads (1.29% of the total number of reads in the library).

## 4 Discussion

High multiplexed approaches for reduced representation libraries such as RAD-seq have considerably reduced the cost of genotyping of hundreds of samples (Bayona-Vásquez *et al*., 2019; Franchini *et al*., 2017). However, library preparation methods introduce a number of artefacts that must be considered when designing a RAD-seq study and its analysis (Andrews *et al*., 2016). The formation of sequences with chimeric adapters, particularly the ones produced by index hopping, is one of such artefacts, which a number of previous studies have attempted to quantify (MacConaill *et al*., 2018; Van Der Valk *et al*., 2019; Costello *et al*., 2018).

Here, we extend these analyses to a much larger (total n = 639) and highly multiplexed experiment (86 to 100 samples multiplexed), such that has now become common in ecological and evolutionary genomics research. Additionally, we consider variation in the library preparation protocol that affect the proportions of chimeric sequences by analysing chimeric sequences formed during both sequencing only and during indexing PCR and amplification during sequencing combined. Finally, we assess the effects of barcode rescue on the proportion of chimeric reads identified and quantify specific types of chimeric sequences (types II-IV) that are impossible to detect in a typical experiment with the same number of samples in every multiplexed group. As our model system *Apodemus spp* does not have the reference genome available, we were not able to identify intra-individual chimeras: chimeric sequences produced between different sequences from the same individual. Such sequences can typically be removed from downstream analysis by mapping to a reference genome.

Overall, we show that the proportion of chimeric sequences is generally low for type A libraries: mean=0.65%, median=0.59%, stdev=0.21. Pooling samples early in the protocol (prior to the indexing PCR, as in our type B libraries) roughly doubles the proportion of detectable chimeric sequences: mean=1.15%, median=1.09%, stdev=0.43, thus increasing read misassignment.

We also show that this proportion is relatively stable throughout several sequencing runs (Supplementary Materials Table 4, 4). In our case, more chimeric sequences identified in library type A-4 have likely arisen due to inclusion of degraded DNA samples. As read length negatively correlates with the frequency of index hopping in a sequencing library (Van Der Valk *et al*., 2019), it could explain the greater proportion of chimeric reads in type A-4 library compared to other runs of libraries of type A.

Previous studies (Van Der Valk *et al*., 2019) have identified similar percentages of chimeric reads - 0.47% - to those obtained in our type A libraries, in a similar protocol that eliminated the possibility of generating chimeras at the indexing PCR stage. In other works, higher proportions of chimeric reads were reported, including in the PCR-free protocols (1.5%, (Ros-Freixedes *et al*., 2018)), Illumina Guidelines (2%, (Illumina, 2017)) or 1.2% in a study by Costello *et al*. (2018). The latter study explained their proportions by low yield of the libraries and a high proportion of free-floating primers on the flow cell.

Our experimental design, with all barcodes separated by 4 nucleotides to minimise read mis-assignment due to sequencing errors, demonstrated that increasing the number of allowed mismatches during barcode rescue in a demultiplexing step increases the proportion of chimeric sequences by as much as 10 fold (Figure 3). Our data shows that allowing more than 2 nucleotides difference in barcode rescue results in extremely high proportion of chimeric sequences, to the point where they become more prevalent in the data than the increase in the number of retained reads due to barcode rescue.

Our results indicate that chimeras type I are the most frequent type of chimeras (0.617% and 1.082% of the total reads for libraries of Type A and Type B respectively), having in mind that our protocol was dominated by equally-sized multiplexed groups. Although the frequency of non-Type I chimeric sequences detected in our protocol is much lower (0.101%, 0.005% and 0.07%, of the total reads for chimeras Type II, III and IV in libraries Type A; 0.112%, 0.02% and 0.072%, of the total reads for chimeras of Type II, III and IV in libraries Type B), most chimeras of Type IV are undetectable in our protocol. If our protocol allowed for detection of all Type IV chimeras, we estimate that they would constitute a 100-fold larger fraction of chimeric sequences.

Therefore, the principal issue with Type IV chimeras is that they are misassigned as genuine samples and they are impossible to detect in the analysis pipeline when multiplexed groups are of the same size. Chimeras Type II and III, in contrast, are routinely classified as chimeras Type I in RAD-seq protocols with equally-sized multiplexed groups and therefore can be removed during the analysis. It is overall, however, difficult to assess the impact of chimeric sequences on downstream analyses and simulations would be needed to estimate the effects of chimeric sequences incorporated into the final genotypes. Nevertheless, we can suggest steps that would minimise their impact in any highly-multiplex RAD-seq experiment.

In experimental designs where costs are less constrained, we would recommend use of fixed pairs of inner and outer adapters for each sample and multiplexed groups, respectively. Although this increases the cost associated with development and adapter synthesis, fixed pairs of barcodes will minimise the probability of biasing downstream analyses due to read misassignment (Van Der Valk *et al*., 2019). We would also recommend performing indexing PCRs on each sample individually. PCR duplicates might have little effect on genotype calls (Euclide *et al*., 2020), however, we still recommend the inclusion of a random nucleotides to identify them. The major benefit of being able to identify PCR duplicates and chimeras is the ability to increase the number of PCR cycles in the samples’ amplification step, increasing the amount of input material available. In cases where the input materials is scarce and/or degraded, controlling for chimeric sequences becomes more important, as their proportion increases with shorter read length and increasing number of mismatches allowed during barcode rescue.

When costs are a limiting factor, we suggest adopting a hybrid approach, similar to the one described here: use of fixed pairs of inner barcodes only and pooling the samples for the indexing PCR. This approach still enables adequate control of the chimeric sequences in the data, while saving costs during library preparation. Nevertheless when using this approach, one should consider not including all combination of inner barcodes in every multiplexed group to be able to estimate the frequency of chimeras type IV that are being misassigned to genuine samples.

## 5 Acknowledgements

The authors wish to thank all collaborators that have contributed samples used in this project (in alphabetical order): Dr Jan Boratyński, Mammal Research Institute, Polish Academy of Sciences, Białowieża, Poland, Dr Dougie Clarke, University of Huddersfield, UK, Dr Jerry Herman, National Museums Scotland, Edinburgh, UK, Dr Vladimir Jovanovic, Freie Universität Berlin, Germany, Dr Johan Michaux, University of Liège, Belgium, Dr Joana Pauperio, CIBIO, University of Porto, Portugal and Dr Karol Zub, Mammal Research Institute Polish Academy of Sciences, Białowieża, Poland. MLMC, RR and JB also wish to acknowledge the use of the Orion High Performance Computing cluster at the University of Huddersfield.

## 6 Data Accessibility

All the code and the demultiplexing information produced by process_radtags, as well as the tables constructed used to perform all the analyses described here are available on GitHub: https://github.com/Marisa89/chimeric_adapters. The data from Type A libraries is available in EBI SRA repository under accession number PRJNA554851. The data from Type B libraries will be made available upon acceptance of the manuscript.

## References

Andrews, K.R., Good, J.M., Miller, M.R., Luikart, G., and Hohenlohe, P.A. (2016). “Harnessing the power of RADseq for ecological and evolutionary genomics”. In: Nature Reviews Genetics 17.2, p. 81.

Baird, Nathan A, Etter Paul D, Atwood Tressa S, Currey Mark C, Shiver Anthony L, Lewis Zachary A, Selker Eric U, Cresko William A, and Johnson Eric A (2008). “Rapid SNP discovery and genetic mapping using sequenced RAD markers”. In: PloS one 3.10, e3376.

Bayona-Vásquez, N.J., Glenn, T.C., Kieran, T.J., Pierson, T.W., Hoffberg, S.L., Scott, P.A., Bentley, K.E., Finger, J.W., Louha, S., Troendle, N., Diaz-Jaimes, P., Mauricio, R., and Faircloth, B.C. (2019). “Adapterama III: Quadruple-indexed, double/triple-enzyme RAD-seq libraries (2RAD/3RAD)”. In: PeerJ 7, e7724.

Catchen, J.M., Amores, A., Hohenlohe, P., Cresko, W., and Postlethwait, J.H. (2011). “Stacks: building and genotyping loci de novo from short-read sequences”. In: G3: Genes, genomes, genetics 1.3, pp. 171–182.

Costello, M., Fleharty, M., Abreu, J., Farjoun, Y.i, Ferriera, S., Holmes, L., Granger, B., Green, L., Howd, T., Mason, T., Vicente, G., Dasilva, M., Brodeur, W., DeSmet, T., Dodge, S., Lennon, N.J., and Gabriel, S. (2018). “Characterization and remediation of sample index swaps by non-redundant dual indexing on massively parallel sequencing platforms”. In: BMC genomics 19.332.

Euclide, Peter T, McKinney Garrett J, Bootsma, Matthew, Tarsa, Charlene, Meek Mariah H, and Larson Wesley A (2020). “Attack of the PCR clones: Rates of clonality have little effect on RAD-seq genotype calls”. In: Molecular ecology resources 20.1, pp. 66–78.

Faircloth, B.C. and Glenn, T.C. (2012). “Not all sequence tags are created equal: designing and validating sequence identification tags robust to indels”. In: PloS one 7.8, e42543.

Fonseca, V.G, Nichols, B., Lallias, D., Quince, C., Carvalho, G.R., Power, D.M., and Creer, S. (2012). “Sample richness and genetic diversity as drivers of chimera formation in nSSU metagenetic analyses”. In: Nucleic Acids Research 40.9, e66–e66.

Franchini, P., Monné Parera, D., Kautt, A.F., and Meyer, A. (2017). “quaddRAD: a new high-multiplexing and PCR duplicate removal ddRAD protocol produces novel evolutionary insights in a nonradiating cichlid lineage”. In: Molecular Ecology 26.10, pp. 2783–2795.

Gao, Y., Yin, S., Wu, L., Dai, D., Wang, H., Liu, C., and Tang, L. (2017). “Genetic diversity and structure of wild and cultivated Amorphophallus paeoniifolius populations in southwestern China as revealed by RAD-seq”. In: Scientific reports 7.1, pp. 1–10.

Glenn, T.C., Nilsen, R.A., Kieran, T.J., Sanders, J.G., Bayona-Vásquez, N. J, Finger, J.W., Pierson, T.W., Bentley, K.E., Hoffberg, S.L., Louha, S., García-De León, F.J., Del Río-Portilla, M.A., Reed, K.D., Anderson, J.L., Meece, J.K., Aggrey, S.E., Rekaya, R., Alabady, M., Bélanger, M., Winker, K., and Faircloth, B.C. (2019). “Adapterama I: universal stubs and primers for 384 unique dual-indexed or 147,456 combinatorially-indexed Illumina libraries (iTru & iNext)”. In: PeerJ 7, e7755.

Hohenlohe, P.A., Day, M.D., Amish, S.J., Miller, M.R., Kamps-Hughes, N., Boyer, M.C., Muhlfeld, C.C., Allendorf, F.W., Johnson, E.A., and Luikart, G. (2013). “Genomic patterns of introgression in rainbow and westslope cutthroat trout illuminated by overlapping paired-end RAD sequencing”. In: Molecular ecology 22.11, pp. 3002–3013.

Illumina (2017). “Effects of index misassignment on multiplexing and downstream analysis”. In: URL: https://www.illumina.com/content/dam/illumina-marketing/documents/products/whitepapers/index-hopping-white-paper-770-2017-004.pdf.

Jeffries, D.L., Copp, G.H., Lawson Handley, L., Olsén, K.H., Sayer, C.D., and Hänfling, B. (2016). “Comparing RADseq and microsatellites to infer complex phylogeographic patterns, an empirical perspective in the Crucian carp, Carassius carassius, L.” In: Molecular ecology 25.13, pp. 2997–3018.

Lecaudey, L.A., Schliewen, U.K, Osinov, A.G., Taylor, E.B., Bernatchez, L., and Weiss, S.J. (2018). “Inferring phylogenetic structure, hybridization and divergence times within Salmoninae (Teleostei: Salmonidae) using RAD-sequencing”. In: Molecular phylogenetics and evolution 124, pp. 82–99.

Leone, A., Álvarez, P., García, D., Saborido-Rey, F., and Rodriguez-Ezpeleta, N. (2019). “Genomewide SNP based population structure in European hake reveals the need for harmonizing biological and management units”. In: ICES Journal of Marine Science 76.7, pp. 2260– 2266.

MacConaill, L.E, Burns, R.T, Nag, A., Coleman, H. A, Slevin, M.K, Giorda, K., Light, M., Lai, K., Jarosz, M., McNeill, M.S, Ducar, M.D., Meyerson, M., and Thorner, A.R. (2018). “Unique, dual-indexed sequencing adapters with UMIs effectively eliminate index cross-talk and significantly improve sensitivity of massively parallel sequencing”. In: BMC genomics 19.30.

Martin Cerezo, M.L, Kucka, M., Zub, K., Chan, Y.F, and Bryk, J. (2020). “Population structure of Apodemus flavicollis and comparison to Apodemus sylvaticus in northern Poland based on RAD-seq”. In: BMC genomics 21.241.

Massatti, R., Reznicek, A. A, and Knowles, L.L. (2016). “Utilizing RADseq data for phylogenetic analysis of challenging taxonomic groups: A case study in Carex sect. Racemosae”. In: American Journal of Botany 103.2, pp. 337–347.

Nadeau, N.J., Ruiz, M., Salazar, P., Counterman, B., Medina, J.A, Ortiz-Zuazaga, H., Morrison, A., McMillan, W.O., Jiggins, C.D, and Papa, R. (2014). “Population genomics of parallel hybrid zones in the mimetic butterflies, H. melpomene and H. erato”. In: Genome research 24.8, pp. 1316–1333.

Near, T.J., MacGuigan, D.J., Parker, E., Struthers, C.D., Jones, C.D., and Dornburg, A. (2018). “Phylogenetic analysis of Antarctic notothenioids illuminates the utility of RADseq for resolving Cenozoic adaptive radiations”. In: Molecular phylogenetics and evolution 129, pp. 268–279.

Peterson, B.K., Weber, J.N., Kay, E.H., Fisher, H.S., and Hoekstra, H.E. (2012). “Double digest RADseq: an inexpensive method for de novo SNP discovery and genotyping in model and non-model species”. In: PloS one 7.5, e37135.

Poland, J.A. and Rife, T.W. (2012). “Genotyping-by-sequencing for plant breeding and genetics”. In: The Plant Genome 5.3, pp. 92–102.

Rodríguez-Ezpeleta, N., Bradbury, I.R., Mendibil, I., Álvarez, P., Cotano, U., and Irigoien, X. (2016). “Population structure of Atlantic mackerel inferred from RAD-seq-derived SNP markers: Effects of sequence clustering parameters and hierarchical SNP selection”. In: Molecular ecology resources 16.4, pp. 991–1001.

Ros-Freixedes, R., Battagin, M., Johnsson, M., Gorjanc, G., Mileham, A.J., Rounsley, S.D., and Hickey, J.M. (2018). “Impact of index hopping and bias towards the reference allele on accuracy of genotype calls from low-coverage sequencing”. In: Genetics Selection Evolution 50.1, p. 64.

Sinha, R., Stanley, G., Gulati, G.S., Ezran, C., Travaglini, K.J., Wei, E., Chan, C.K.F., Nabhan, A.N., Su, T., Morganti, R.M., Conley, S.D., Chaib, H., Red-horse, K., Longaker, M.T., Snyder, M.P., and Krasnow M.A. abd Weissman, I.L. (2017). “Index switching causes “spreading-of-signal” among multiplexed samples in Illumina HiSeq 4000 DNA sequencing”. In: BioRxiv, p. 125724.

Smyth, R.P, Schlub, T.E, Grimm, A, Venturi, V., Chopra, A., Mallal, S., Davenport, M.P, and Mak, J. (2010). “Reducing chimera formation during PCR amplification to ensure accurate genotyping”. In: Gene 469.1-2, pp. 45–51.

Van Der Valk, T., Vezzi, F., Ormestad, M., Dalen, L., and Guschanski, K. (2019). “Index hopping on the Illumina HiseqX platform and its consequences for ancient DNA studies”. In: Molecular ecology resources.

Wang, G.C.Y. and Wang, Y. (1996). “The frequency of chimeric molecules as a consequence of PCR co-amplification of 16S rRNA genes from different bacterial species”. In: Microbiology 142.5, pp. 1107–1114.

