## Supplementary materials for "Identification and quantification of chimeric sequencing reads in a highly multiplexed RAD-seq protocol"

Table 1: i5 adapter's sequences

| **Adapter name** | **Sequence** |
|---|---|
| i5-top_#01_AAGACTGG | CGCTCTTCCGATCTVBBNAAGACTGGTGCA/3Phos/ |
| i5-top_#02_ATGTTGGC | CGCTCTTCCGATCTVBBNATGTTGGCTGCA/3Phos/ |
| i5-top_#04_CCTCATCT | CGCTCTTCCGATCTVBBNCCTCATCTTGCA/3Phos/ |
| i5-top_#05_CGGAATTG | CGCTCTTCCGATCTVBBNCGGAATTGTGCA/3Phos/ |
| i5-top_#06_CAAGGTGA | CGCTCTTCCGATCTVBBNCAAGGTGATGCA/3Phos/ |
| i5-top_#07_GACTTGAG | CGCTCTTCCGATCTVBBNGACTTGAGTGCA/3Phos/ |
| i5-top_#10_TCCTTCAC | CGCTCTTCCGATCTVBBNTCCTTCACTGCA/3Phos/ |
| i5-top_#11_TGTCAGTG | CGCTCTTCCGATCTVBBNTGTCAGTGTGCA/3Phos/ |
| i5-top_#12_TTCTGAGG | CGCTCTTCCGATCTVBBNTTCTGAGGTGCA/3Phos/ |
| i5-bottom_#01_AAGACTGG | /5Phos/CCAGTCTTNVVBAGATCGGAAGAGCGTCGTGTAGGGAAAGAGTGT |
| i5-bottom_#02_ATGTTGGC | /5Phos/GCCAACATNVVBAGATCGGAAGAGCGTCGTGTAGGGAAAGAGTGT |
| i5-bottom_#04_CCTCATCT | /5Phos/AGATGAGGNVVBAGATCGGAAGAGCGTCGTGTAGGGAAAGAGTGT |
| i5-bottom_#05_CGGAATTG | /5Phos/CAATTCCGNVVBAGATCGGAAGAGCGTCGTGTAGGGAAAGAGTGT |
| i5-bottom_#06_CAAGGTGA | /5Phos/TCACCTTGNVVBAGATCGGAAGAGCGTCGTGTAGGGAAAGAGTGT |
| i5-bottom_#07_GACTTGAG | /5Phos/CTCAAGTCNVVBAGATCGGAAGAGCGTCGTGTAGGGAAAGAGTGT |
| i5-bottom_#10_TCCTTCAC | /5Phos/GTGAAGGANVVBAGATCGGAAGAGCGTCGTGTAGGGAAAGAGTGT |
| i5-bottom_#11_TGTCAGTG | /5Phos/CACTGACANVVBAGATCGGAAGAGCGTCGTGTAGGGAAAGAGTGT |
| i5-bottom_#12_TTCTGAGG | /5Phos/CCTCAGAANVVBAGATCGGAAGAGCGTCGTGTAGGGAAAGAGTGT |

Table 2: i7 adapter's sequences

| **Adapter name** | **Sequence** |
|---|---|
| i7-top_#01_AGAGTTCG | GTGACTGGAGTTCAGACGTGTGCTCTTCCGATCTVBBNAGAGTTCG |
| i7-top_#02_ACCTGTTG | GTGACTGGAGTTCAGACGTGTGCTCTTCCGATCTVBBNACCTGTTG |
| i7-top_#03_CTGGTTCA | GTGACTGGAGTTCAGACGTGTGCTCTTCCGATCTVBBNCTGGTTCA |
| i7-top_#04_CGACAAGA | GTGACTGGAGTTCAGACGTGTGCTCTTCCGATCTVBBNCGACAAGA |
| i7-top_#05_CAGTCGAA | GTGACTGGAGTTCAGACGTGTGCTCTTCCGATCTVBBNCAGTCGAA |
| i7-top_#06_GTCAGAAC | GTGACTGGAGTTCAGACGTGTGCTCTTCCGATCTVBBNGTCAGAAC |
| i7-top_#7_TTGTTCCG | GTGACTGGAGTTCAGACGTGTGCTCTTCCGATCTVBBNTTGTTCCG |
| i7-top_#8_TCGCATTC | GTGACTGGAGTTCAGACGTGTGCTCTTCCGATCTVBBNTCGCATTC |
| i7-top_#9_TCGAACCA | GTGACTGGAGTTCAGACGTGTGCTCTTCCGATCTVBBNTCGAACCA |
| i7-bottom_#01_AGAGTTCG | TACGAACTCTNVVBAGATCGGAAGAGCA |
| i7-bottom_#02_ACCTGTTG | TACAACAGGTNVVBAGATCGGAAGAGCA |
| i7-bottom_#03_CTGGTTCA | TATGAACCAGNVVBAGATCGGAAGAGCA |
| i7-bottom_#04_CGACAAGA | TATCTTGTCGNVVBAGATCGGAAGAGCA |
| i7-bottom_#05_CAGTCGAA | TATTCGACTGNVVBAGATCGGAAGAGCA |
| i7-bottom_#06_GTCAGAAC | TAGTTCTGACNVVBAGATCGGAAGAGCA |
| i7-bottom_#7_TTGTTCCG | TACGGAACAANVVBAGATCGGAAGAGCA |
| i7-bottom_#8_TCGCATTC | TAGAATGCGANVVBAGATCGGAAGAGCA |
| i7-bottom_#9_TCGAACCA | TATGGTTCGANVVBAGATCGGAAGAGCA |

Table 3: Combinatorial outer adapter sequences

| **Adapter name** | **Sequence** |
|---|---|
| i501_AGCATGGA | AATGATACGGCGACCACCGAGATCTACAC{AGCATGGA}ACACTCTTTCCCTACACGAC*G |
| i502_CCTGGAAT | AATGATACGGCGACCACCGAGATCTACAC{CCTGGAAT}ACACTCTTTCCCTACACGAC*G |
| i503_GCAAGCAA | AATGATACGGCGACCACCGAGATCTACAC{GCAAGCAA}ACACTCTTTCCCTACACGAC*G |
| i504_TGAGGATG | AATGATACGGCGACCACCGAGATCTACAC{TGAGGATG}ACACTCTTTCCCTACACGAC*G |
| i701_ACACTCAG | CAAGCAGAAGACGGCATACGAGAT{CTGAGTGT}GTGACTGGAGTTCAGACGTGTGC*T |
| i702_CAGTCGAA | CAAGCAGAAGACGGCATACGAGAT{TTCGACTG}GTGACTGGAGTTCAGACGTGTGC*T |
| i703_GGCTCAAT | CAAGCAGAAGACGGCATACGAGAT{ATTGAGCC}GTGACTGGAGTTCAGACGTGTGC*T |
| i704_TTCCGCTT | CAAGCAGAAGACGGCATACGAGAT{AAGCGGAA}GTGACTGGAGTTCAGACGTGTGC*T |

Table 4: Proportion of chimeric sequences per plate. Median (+/- standard deviation), mean, maximum and minimim values are shown.

| Plate | Mismatches | Median | Stdev | Mean | Max | Min |
|---|---|---|---|---|---|---|
| PlateA-1 | 0.00 | 0.54 | 0.14 | 0.55 | 0.87 | 0.36 |
| PlateA-2 | 0.00 | 0.59 | 0.17 | 0.64 | 0.91 | 0.38 |
| PlateA-3 | 0.00 | 0.49 | 0.12 | 0.53 | 0.69 | 0.33 |
| PlateA-4 | 0.00 | 0.90 | 0.23 | 0.89 | 1.21 | 0.44 |
| PLateB-1 | 0.00 | 0.83 | 0.19 | 0.89 | 1.39 | 0.68 |
| PlateB-2 | 0.00 | 1.32 | 0.16 | 1.30 | 1.51 | 1.03 |
| PLateB-3 | 0.00 | 1.25 | 0.67 | 1.28 | 2.33 | 0.31 |
